# Radiation Threats to Humans in Space and an alternative approach with Probiotics

**DOI:** 10.1101/2021.03.12.435153

**Authors:** Dilara Diken, Elia Brodsky, Harpreet Kaur, Mohit Mazumder

## Abstract

Space type radiation is an important factor to consider for scientists on International Space Stations, especially high linear transfer energy (LET) since it has imminent effects on microorganisms. The abundances of bacteria are a good indicator of how radiation influences the gut microbiome. The current study is an attempt towards this; thus, we have employed a public dataset (Bioproject code PRJNA368790) of 80 mice samples treated with a range of doses from 0Gy to 1Gy and feces samples were collected at different time points of post radiation treatment. Metagenomic analysis was performed on this data to understand the effect of radiation doses on the abundance of microbial species or microbial diversity implementing the DADA2 and Phyloseq pipelines. Our analyses have shown that 0.1Gy high LET radiation had the significant effect on the species of bacteria. There is a significant decrease in four types of bacteria, i.e., Bifidobacterium longum, Bifidobacterium castoris, Lactobacillus gasseri and Lactobacillus johnsonii, with p-value of 7.05×10-5, 0.020, 0.057and 0.020, respectively. Additionally, pathway analysis indicates the protein coding products of these bacteria are involved in the GABAergic synaptic pathway. Further, our study has shown the significant difference between post radiation time points, i.e., 10 days vs.30 days and suggested the acclimatization period could be around 10 days for these bacteria.

## INTRODUCTION

Radiation in general is categorized into ionizing and non-ionizing radiation. Ionizing radiation is a greater threat than non-ionizing radiation due to its chemical properties; when it passes through substances it is able to cause chemical reactions since it has high energy (Perez, 2017). High linear energy transfer (LET), which is a type of ionizing radiation, causes damage on several locations along the DNA that are close together (Takahashi, Ikeda, & Yoshida, 2018). The effects of radiation on somatic cells extends to micro-organisms, too (Maalouf, Durante, & Foray, 2011). These researchers have also stated that it was difficult to draw conclusions from the data available due to different metabolic rates of individuals. Therefore, more data would be required to conclusively suggest a relationship between some conditions under study and radiation. Numerous mechanisms of microbes are affected by radiation. These mechanisms include growth, gene expression and repair; the effects are not limited to microbial survival but also host-microbe relations (Senatore et al., 2018).

Although we are somewhat shielded from space-type radiation on Earth; however, scientists and astronauts on International Space Stations (ISS) are exposed to higher doses. Astro-scientists have conducted over a thousand studies on space stations ranging from biology to physics (Witze, 2020). The health and well-being of these scientists are of utmost importance to be able to continue this research. According to reports from NASA (2015) where they discuss issues that are largely uncontrolled, partially controlled, and completely controlled, radiation is and will continue to be uncontrolled for the foreseeable future. There are physical shields or force fields used to shield from galactic cosmic rays (Garner, 2015), but more creative methods might be required to decrease the relative biological effects of radiation.

Bacteria was commonly studied individually until the rise of NGS; this novel sequencing technology allows us to study communities of bacteria in a sample (Maccaferri, Biagi, &Brigidi, 2011), such as the microbiome. Gut microbiome is affected by many external factors ranging from sleep disorder to psychological stress (Lobionda, Sittipo, Kwon, & Lee, 2019). On the other hand, dysbiosis-a decrease in good gut microbes, triggers a decrease in chemicals necessary for the nervous system, which eventually leads to psychological disorders (Eltokhi, Janmaat, Genedi, Haarman, & Sommer, 2020). Hence there is a cyclic nature in the progression of psychological disorders. The gut-brain axis also refers to the common saying your gut is your second brain. This connection is possible due to the vagus nerve that connects the central nervous system and the intestines by interacting with the entero-dendritic nerve (Bonaz, Bazin, & Pellissier, 2018). This systemic connection is more feasible than a connection through the circulatory system since the blood-brain is extremely partial to chemicals.

Moreover, metabolites absorbed in the gut can lead to divergence on behavior because of the evolutionary symbiotic relationship between bacteria and their hosts. Micro-organisms prosper by tweaking their hosts natural functions to better suit their needs (Johnson & Foster, 2018). Johnson & Foster (2018) also state that bacterial species in the Lactobacillus and Bifidobacterium species affect the nervous system and hence psychology through metabolites like GABA. Gama aminobutyric acid is known to be the chief inhibitory neurotransmitter in the CNS, it is produced by the enzyme GAD (glutamate decarboxylase) (Duman, Sanacora, & Krystal, 2019). A lack of GABA or the decreased stimulation of GABA receptors causes neurobehavioral disorders like depression and anxiety (Kalueff & Nutt, 2007). There are several treatment options available to treat conditions that arise due to dysbiosis. One course of action is through fecal transplantations (Holleran et al., 2018). On the other hand, some researchers use probiotics to overcome these afflictions (Tabouy et al., 2018), which is a simple, cost effective and non-invasive method.

As aforementioned radiation has an effect on microorganisms and to study the complications that astronauts face due to the extreme conditions in space scientists are developing new techniques. NASA has a radiation center to facilitate research to better understand the short- and long-term effects on model organisms like mice (Tessa, Sivertz, Chiang, Lowenstein, & Rusek, 2016). In this article data from previous studies from NCBI was used to test if the properties of high LET radiation are true and if the abundances of bacteria decrease significantly.

## METHODS

### Dataset & Experimental Design

The dataset was obtained from the Bioprojects section on NCBI under the article named “Space-type radiation induces multimodal responses in the mouse gut microbiome and metabolome” with the accession code PRJNA368790 (Casero et al., 2017). The mice in this experiment were exposed to 0.1Gy, 0.25Gy and 1Gy radiation at the NASA Space Radiation Laboratory. The control group were not exposed to radiation but they were kept under the same housing and nutritional conditions. Each of the groups consist of 10 male mice at 6 months old for reproducibility. The gut microbiome was the focus of this study therefore fecal samples were collected from the mice at 10 days and 30 days post radiation.

### Pipeline

For metagenomic analysis of paired end 16S rRNA the DADA2 pipeline is recommended since it can identify between species even with just one nucleotide difference (Callahan et al., 2016). T-Bi-oInfo server (Tauber Bioinformatics: Making sense of big data in BIOINFORMATICS (t-bio.info) was used to analyze the samples obtained. As shown in Figure 3 DADA2 was used to calculate the operational taxonomic units (OTU); whereas PHYLOSEQ was used for other quantitative analyses like Shannon Index. The parameters set for expected number of reads forwards and backwards was 150. The expected errors were estimated as 2 and the min/max value for overlaps was approximated as 12/5. The mean abundance was established as 150 after optimization. Once the OTU table was generated the results were scanned for changes in diversity.

### Data Exploration

The data exploration conducted included non-metric multidimensional scaling (NMDS), principal coordinate analysis (PCoA) and principal component analysis (PCA), which are data reduction techniques used to visualize data that is multidimensional on a two-dimensional scale. After obtaining the PHYLOSEQ outputs that contained PCoA and NMDS, PCA plots were generated using the T-BioInfo server (Tauber Bioinformatics: Making sense of big data in BIOINFORMATICS (t-bio.info)) to visualize the separation between groups. Shannon Index plot is a product of PHYLOSEQ, too. From these indices ENS-effective number of species, was calculated; this number indicates a comparable richness and evenness value. Subsequently this index was used to compare alpha diversity.

Another step used for confirmation was clustering. The type of clustering used was hierarchical clustering (H-clustering), which groups the data depending on how closely related they are. This graphic was also generated using the T-BioInfo server.

### Statistical Tests

To confirm that the observations from the PHYLOSEQ pipeline output ANOVA and T.test was calculated in excel. ANOVA also known as the F-test is a technique used to compare groups with more than one category. If F value is greater than F-critical then it suggests that there is a significant difference between the classes being tested. The p-value is another indicator of ANOVA that can be used to signify significance. There are three hypotheses under in question. The first hypothesis (H0) compares the days to check for differences between samples collected 10 days and 30 days post radiation. H1 is the second hypothesis which compares the doses. The doses chosen were 0Gy, 0.1Gy, and 0.25Gy since in the exploratory phase lower doses were most separate. As for the third and final hypothesis (H2) the interaction between doses and days is compared. This test was concentrated on Lactobacillus and Bifidobacterium species after filtering out species with 0 abundance across all groups. The threshold was set as 0.05 for this experiment and as can be seen from Table 1 the general trend indicates that dosages play an important role.

**Table 1.** Dataset (PRJNA368790) representing the number of samples in each group.

|  | 0Gy | 0.1Gy | 0.25Gy | 1Gy |
| --- | --- | --- | --- | --- |
| 10 days | 10 | 10 | 10 | 10 |
| 30 days | 10 | 10 | 10 | 10 |

Since three different doses were tested in H1 to refine the research further and examine which dose had greater significance when compared to 0Gy t.test is chosen. For the t.test only one category can be compared between two groups therefore the four selected species of bacteria were separated by keeping the days constant and testing 0Gy against 0.1Gy and 0.25Gy separately. As the table 2 above indicates that with threshold *p<0.05 10 days post radiation at 0.1Gy radiation is the group that is significantly different from 0Gy.

**Table 2.** ANOVA F-test results representing significance of the effect of radiation doses and time points for the selected four species of bacteria.

| Species | H0 | H1 | H2 |
| --- | --- | --- | --- |
| Lactobacillus gasseri | F-test | F-test | F-test |
|  | 2.05<4.01 | 5.95>3.17 | 5.13>3.17 |
|  | P-value | P-value | P-value |
|  | 0.150 | 0.005 | 0.009 |
| Lactobacillus johnsonii | F-test | F-test | F-test |
|  | 2.17<4.02 | 9.05>3.17 | 1.08<3.17 |
|  | P-value | P-value | P-value |
|  | 0.147 | 0.0004 | 0.344 |
| Bifidobacterium longum | F-test | F-test | F-test |
|  | 10.14>4.02 | 4.73>3.17 | 0.83<3.17 |
|  | P-value | P-value | P-value |
|  | 0.002 | 0.013 | 0.440 |
| Bifidobacterium castoris | F-test | F-test | F-test |
|  | 5.32>4.02 | 4.37>3.17 | 0.05<3.17 |
|  | P-value | P-value | P-value |
|  | 0.025 | 0.017 | 0.95 |

**Table 3.** Results from T.test to compare the doses only by keeping days constant.

| Species |  | 10 days | 30 days |
| --- | --- | --- | --- |
| Lactobacillus gasseri |  |  |  |
|  | 0Gy v. 0.1 Gy | 0.057 | 0.627 |
|  | 0Gy v. 0.25 Gy | 0.083 | 0.511 |
| Lactobacillus johnsonii |  |  |  |
|  | 0Gy v. 0.1 Gy | 0.020 | 0.030 |
|  | 0Gy v. 0.25 Gy | 0.756 | 0.016 |
| Bifidobacterium longum |  |  |  |
|  | 0Gy v. 0.1 Gy | 7.056x10<sup>-5</sup> | 0.300 |
|  | 0Gy v. 0.25 Gy | 0.318 | 0.253 |
| Bifidobacterium castoris |  |  |  |
|  | 0Gy v. 0.1 Gy | 0.020 | 0.266 |
|  | 0Gy v. 0.25 Gy | 0.035 | 0.037 |

### KEGG Analysis

Kyoto Encyclopedia of Genes and Genomes (KEGG) is a database that presents a collection of the genes and cellular pathways in organisms (https://www.genome.jp/kegg/). On the other hand, the IMGM (Integrated Microbial Genomes and Microbiomes) section of the JGI (Joint Genome Institute) is a database that contains annotations of genes available in different species of bacteria (https://img.jgi.doe.gov/). It even connects the genes of bacteria to cellular pathways in their hosts through KEGG, which is why *L.gasseri, L.johnsonii, B.longum, B.castoris* became the focus of this research. These bacteria have protein coding genes in the GABAergic pathway.

## RESULTS

### Exploratory Analyses

The exploratory anlayses within which NMDS, PCoA and PCA was conducted revealed that samples radiated with 0.1Gy high LET radiation and the control group were relatively distinguished from each other (Figure 2a). In this stage the days do not differentiate strongly. Further, the output from the PHYLOSEQ pipeline also revealed that the overall diversity of the mouse gut microbiome decreases for doses 0.1Gy (Figure 2b). Effective number of species (ENS) was calculated from the Shannon Index and the results were compared on the basis of days. The outcome illustrates a decline of about one species in every group except for 0.1Gy. When samples from mice radiated at 0.1Gy were collected at 10 days post radiation there were two species less than the samples that were collected at 30 days post radiation.

**Figure 1.**
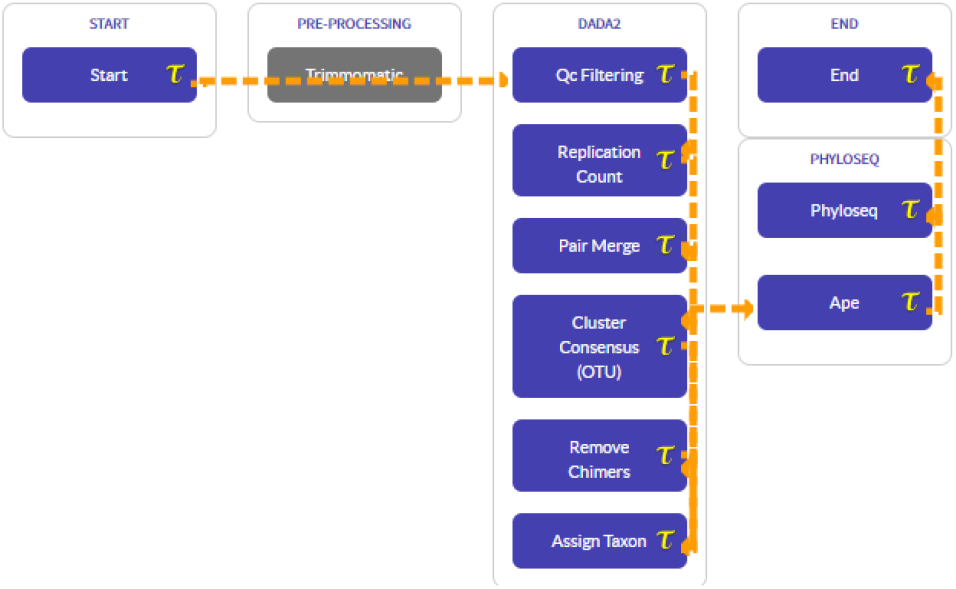
The pipeline used for metagenomics analysis of the gut microbiome of mice that were exposed to radiation. This study utilized both DADA2 and PHYLOSEQ from the T-BioInfo server.

**Figure 2.**
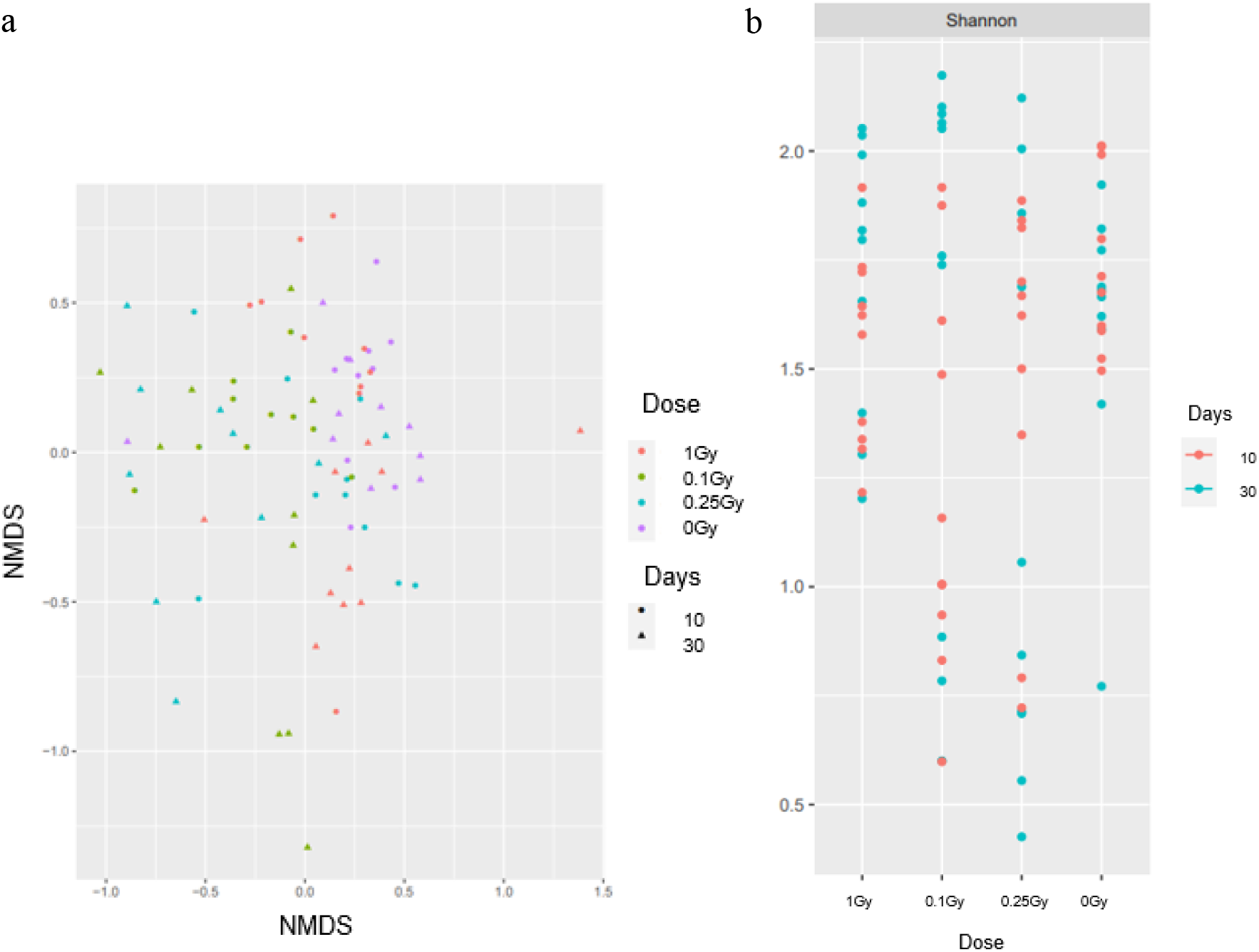
**(a)** Non-metric multidimensional scaling (NMDS) for radiation treated samples. **(b)** The Shannon Index for the four different doses of radiation.

### Phylogenetic Analysis

Furthermore, another output of the PHYLOSEQ pipeline is the phylogenetic tree as given in Figure 3. It shows the relationship of the operational taxonomic units (OTUs) that have significance, and at the same time compare the abundances of OTUs with different dosages and days. From this graphical representation it was inferred that for 10 days post radiation at lower dosages of high LET radiation (0.1Gy) there is lower abundances, especially for OTU 3, 4, and 5.

**Figure 3.**
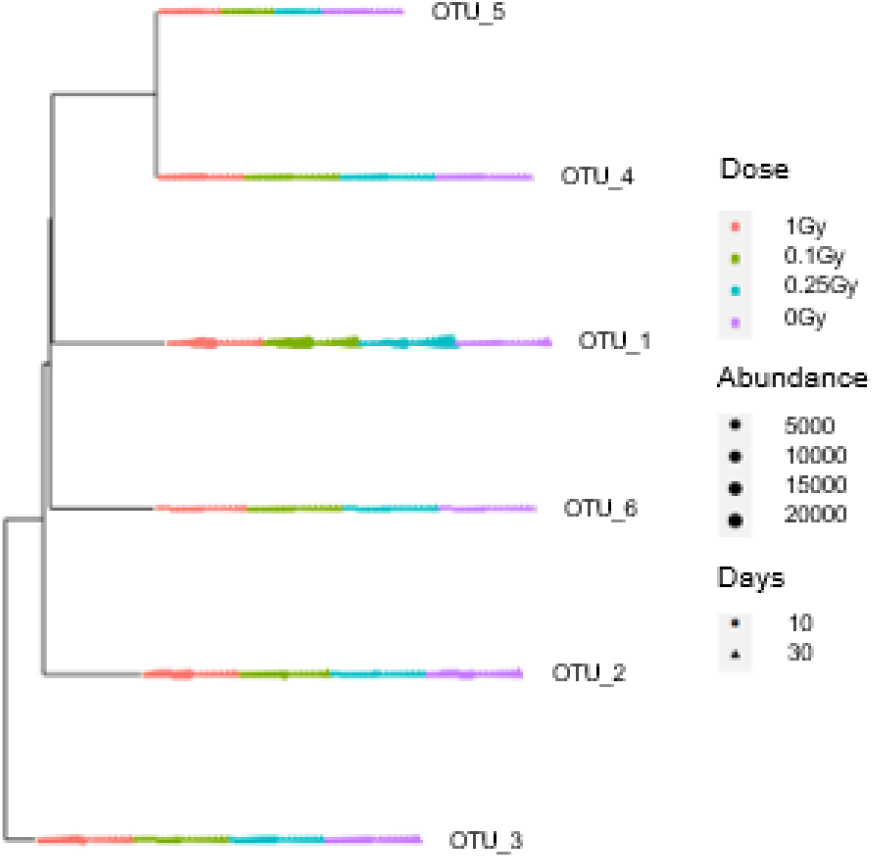
The phylogenetic tree displaying the significant OTUs with their respective abundances correlating with days and dosages.

### Clustering

Next to confirm the observation from the previous analyses, H-clustering was conducted on samples radiated with 0.1Gy and collected at 10 days post radiation. H-clustering was performed to ensure whether samples that belong to one class get clustered together or not. Dendrogram (Figure 4) suggests that there is almost an absolute separation but there are some intermixed samples which can be attributed to noise. Four of each of the samples are mixed in the far-right clade, whereas the rest are distributed in distinctly separate clades.

**Figure 4.**
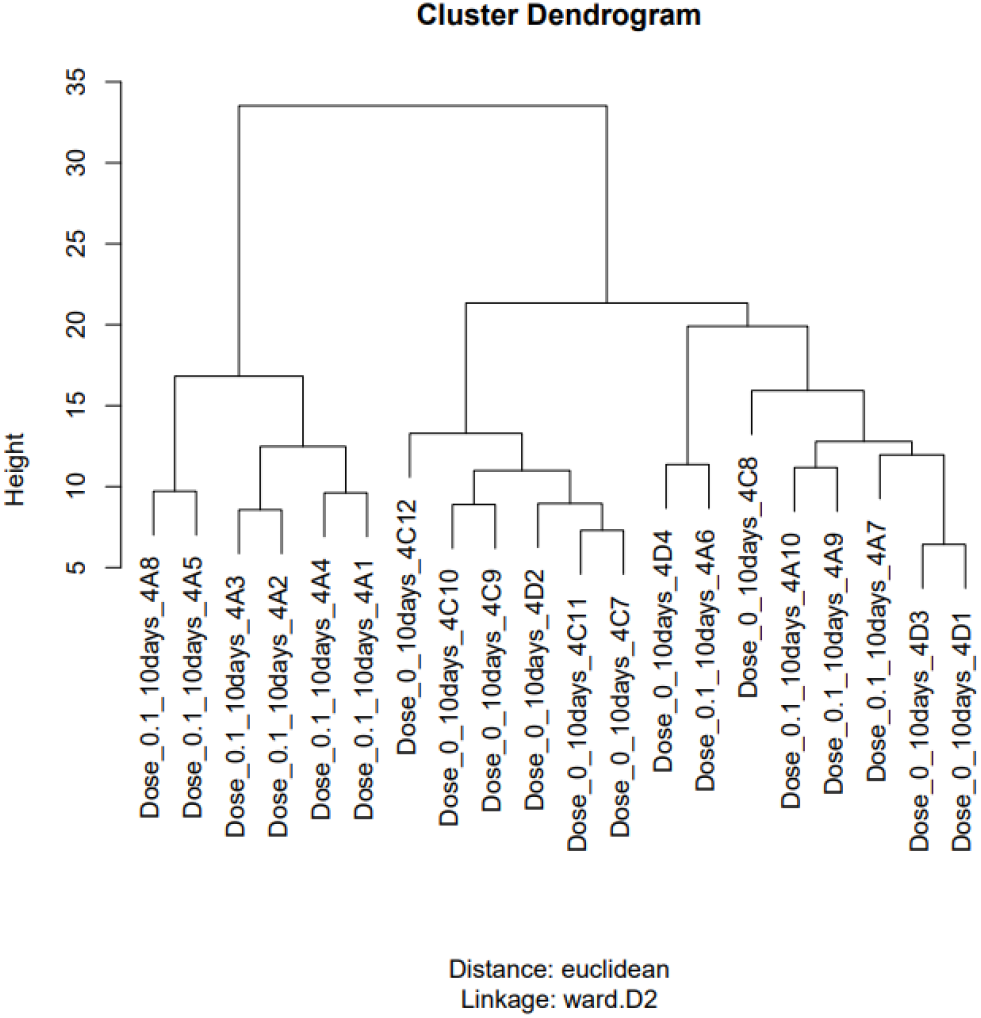
H-cluster diagram for Dose 0Gy and 0.1Gy at 10 days post radiation.

### Statistical Analysis

In addition, statistical tests were conducted after eliminating 1Gy radiation groups. Through ANOVA it was concluded that doses had a greater significance than days or the interaction between categories. T.test was used to decipher which dosage was most significant and 0Gy vs. 0.1Gy displayed a general trend. As the heatmap below suggests (Figure 5) OTU 3, 4, 5, and 8 have decreasing abundances when exposed to 0.1Gy radiation in relative comparison to 0Gy. It is important to note that this figure is only for groups with feces collected at 10 days post radiation. Some of the OTUs are identified as outliers since within the 10 samples there is no separation into two groups as expected. Instead, there is only one or two samples that is different from the others for example, OTU 10 and 12. On the other hand, one OTU that is an exception to the decrement rule is OTU 1, which increases when exposed to 0.1Gy radiation.

**Figure 5.**
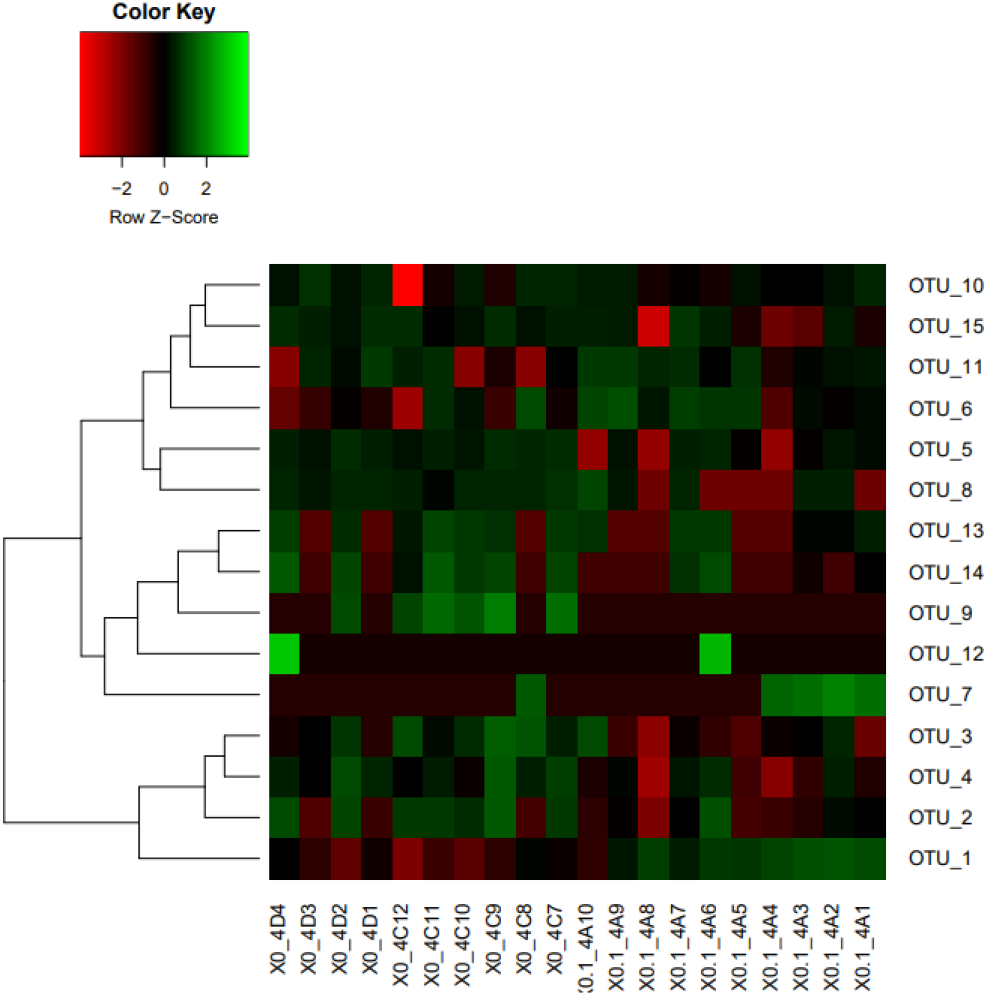
Heatmap representing the OTU abundances in mice that are radiated with dose of 0.1Gy and 0Gy radiation at 10 days.

### OTU Classification

Moreover, another fundamental step that was executed in this research was to identify which species the aforementioned four OTUs belong to. The bar plot below (Figure 6) displays the increase and decrease of abundances of different families of bacteria. Erysipelotrichaceae has the greatest abundance which is on average 30 units and decreases with high LET radiation by 10 units. The second highest abundance was observed in the Lactobacillaceae and Bifidobacteriaceace family 16 units and 12 units respectively. Lactobacillaceae decreased by 7 units whereas Bifidobacte-riaceace decreased by 4 units when exposed to 0.1Gy radiation. These two taxa decrease, whereas Verrucomicrobiacaea follows the opposite trend with a significant increase from 5 units to 23 units when exposed to low dose radiation. The rest of the families are approximately stable under both conditions.

**Figure 6.**
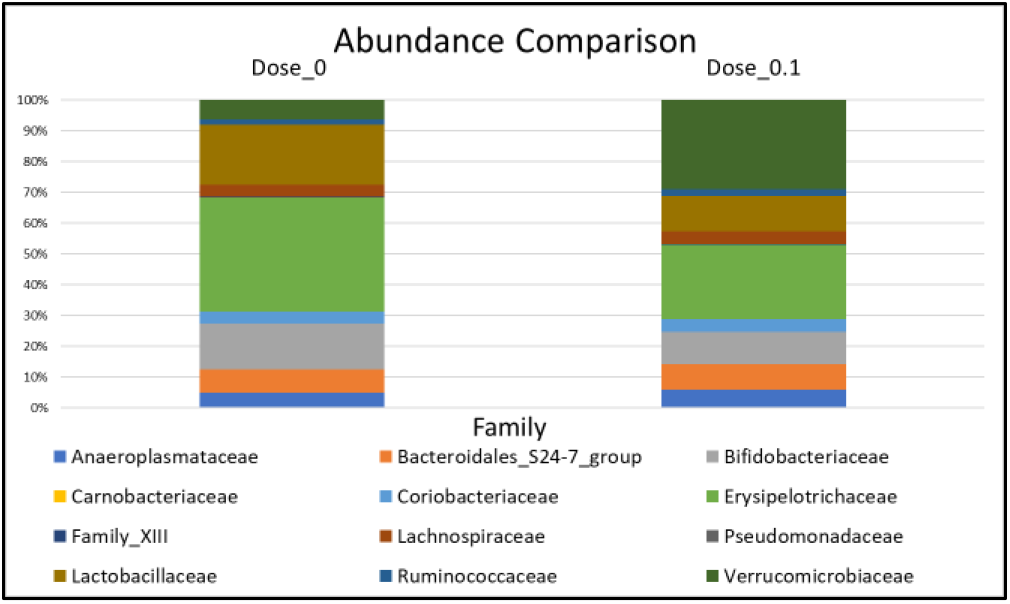
Bar graphs representing the abundances of different families of bacteria in mice that were not radiated and that were radiated at 0.1Gy high LET radiation.

Aligned with previous findings from the PHYLOSEQ pipeline, the significant OTUs are 3, 4, 5, and 8. With further evidence backing up the significant change in the abundances of these four OTUs, their genus and species were identified sing the JGI/IMGM database as follows: OTU 3 is *L.gasseri*, OTU 8 is *L.johnsonii*, OTU 4 is *B.longum* and OTU 5 is *B.castoris*. These bacteria had numerous protein coding genes in several pathways but the relevance of these species in this research is that they have protein products that have an effect on the GABAergic synapse as Figure 7 presents.

**Figure 7.**
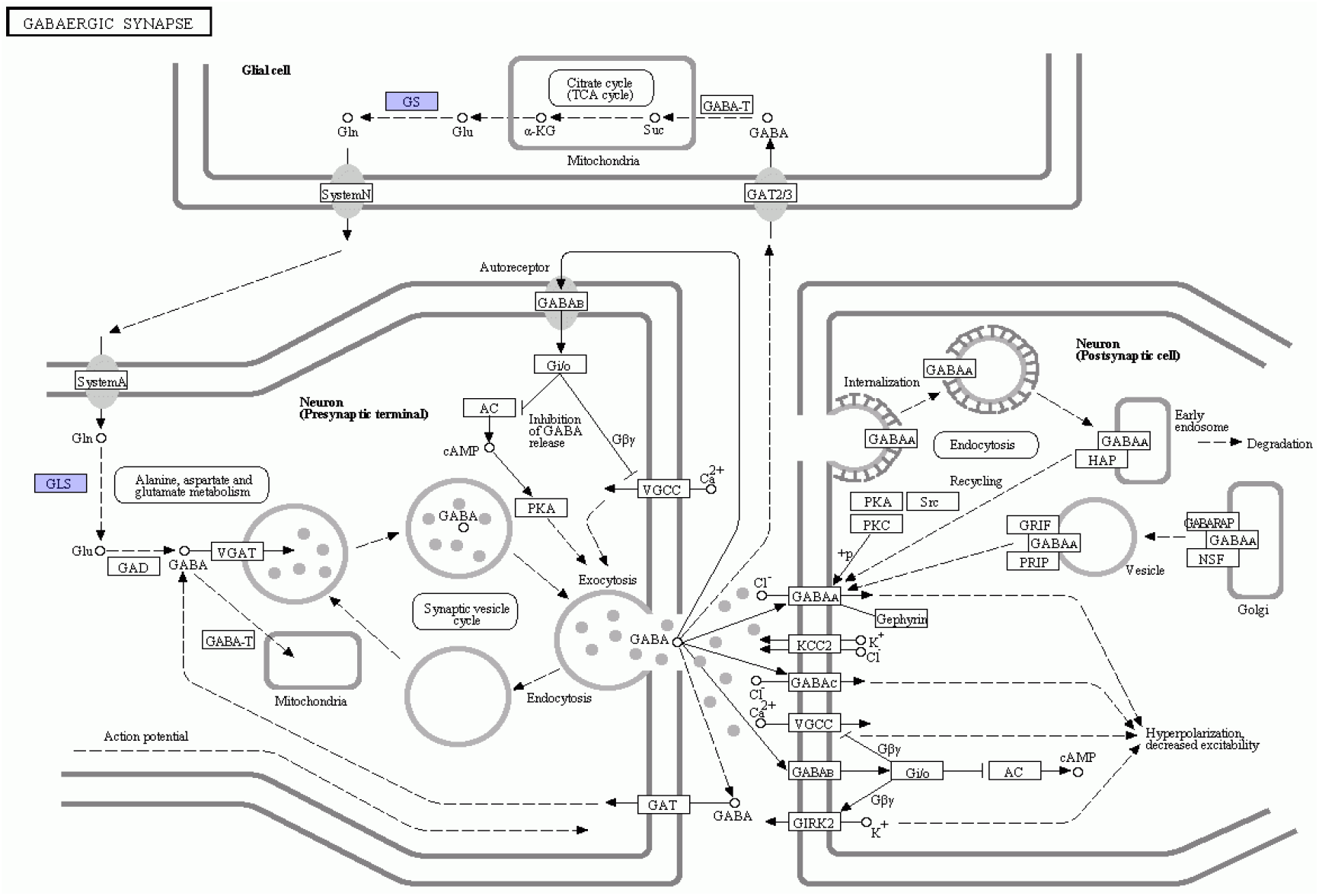
Screenshot from the KEGG database presenting the GABAergic synaptic pathway in which *L.gasseri, L.johnsonii, B.longum* and *B.castoris* have protein coding genes.

The species in the Lactobacillus genus code for proteins involved in both glutaminase (GLS) and glutamine synthase (GS) pathways in the nervous stem. On the other hand, the OTUs in the Bifidobacteria genus were engaged in only one of the pathways in the GABAergic synaptic pathway, which is glutamine synthase (GS). These are both enzymes that process glutamine before it is converted to the neurotransmitter GABA.

## DISCUSSION

The aim for this research was to analyze the effects of high LET radiation on the mouse gut microflora. Th dataset which consisted of 80 samples that was obtained from NCBI was assessed through various steps. Initially, the abundance table was generated using the DADA2 pipeline in the T-Bioinfo server. Complementarily, Phyloseq pipeline was run to produce graphics and estimations to guide the rest of the study. In the following assays NMDS, H-clustering, ANOVA and t.test were used, which consequently condensed the studies’ focus on lower doses. Eventually, the OTUs with significance were identified and these species were connected through nervous system pathways.

From the research we conclude that lower doses particularly 0.1Gy radiation is significantly different than the control group, which did not receive radiation treatment. This result can be attributed to the property of high LET radiation. As the report from ICRU (1970) suggests the relationship between dose and relative biological effect (RBE) is different for different cell types and generally increases for lower doses and then enters a decline state.

Moreover, there is a recurring significance at 10 days post radiation; presumably pointing to the acclimatization period because the collective results signify a return to normal at 30 days post radiation. Since the samples were collected only at two time periods, these results could vary if a more extensive study was conducted. However, this is also linked with the duration of radiation exposure. Mice under study were not exposed to durations that astronauts in deep space travel experience. Therefore, we can call attention to length of exposure as a further limitation that constricts the validity of the results.

The study in question is an important research because the amount of radiation received by astronauts in space travel for almost a year is equivalent to an individual on earth getting a CT scan every week (Tate, 2013). In consideration of the new age goals of corporations like SpaceX, space travel and space tourism will be an important segment of our lives in the upcoming decade (Sheetz, 2020). In light of these developments similar studies should be encouraged. Albeit, this study is only a precursor and was conducted on mice rather than humans, but the bacteria that were specified during this research inhabit humans as host too (Wardeh, Risley, Mcintyre, Setzkorn, & Baylis, 2015).

The main goal of this research was to locate a change that can be connected to neurobehavioral disorders, through observable increments or decrements in the abundances with radiation exposure. Since these bacteria depend on each other in the community the increase of opportunistic bacteria could be due to the decrease of symbiotic bacteria and hence a lack of competition. Thus, an increase in bacteria that produce chemicals that positively affect the host could overcome several conditions caused by imbalance (Gayathri & Rashmi, 2017).

According to studies the two genus of bacteria play a crucial role in the progression of psychological disorders like anxiety and depression (Yunes et al., 2019). GABA which is the most prominent inhibitory neurotransmitter in the CNS will slow down the transmissions in the neurons; this results in a relaxed state. A deficit of this chemical or a lack of stimulation of GABA receptors leads to a neurobehavioral divergence. We conclude that this imbalance in the gut microbiome was the prominent reason as to why staff in the ISS experience anxiety and depression. This trend does not discriminate between different International Space Stations.

A feasible and effective way of increasing stimulation of these receptors that were identified as targets is by increasing bacteria that through the enteric neurons and the vagus nerve activate pathways in the CNS (Bravo et al., 2011). This replicates a neurotypical environment and aids in the treatment of patients. For astronauts in space who have external stressors like increased exposure to radiation, lack of mobility and the extremely sterile environments of their living conditions, it is difficult to propose a practical solution but probiotics seem to be the most efficient course of action.

A continuation to this study could be a cojoined research to study the effects of radiation on psychology by testing mice behavior and try probiotic treatment options. The validity of results will be affirmed further if factors like microgravity are included in future research.

## CONCLUSION

This article has successfully addressed the question of how radiation affects the abundances of bacteria. It is important to note that 0.1Gy radiation had the greatest effect on the species of bacteria that were targeted in this study proving that high LET radiation has greater penetrative qualities at lower dosages for microorganisms. Additionally, 10 days post radiation stood out in comparison to 30 days post radiation; this indicates that the acclimatization period could be around 10 days. Bacteria in Lactobacillus and Bifidobacterium taxon decreased significantly, which can be connected to the decrease in GABA. Since these bacteria have protein coding genes that affect the nervous system, we concluded that astronauts afflicted by anxiety and depression could also be suffering from dysbiosis. Hence a recommendation of probiotics rich in lactobacillus and bifidobacteria is fundamental.

These studies should be extended to testing on fecal samples of astronauts who have been on long duration space travel in order for a better understanding of how problems arising from the extreme conditions in space and the ISS can be overcome. Besides in reality high LET radiation is combined with other types of radiation, microgravity and high sterility, therefore increasing the complexity of both the problem and the solution. If space tourism is to be a staple of the future it is important to continue similar research.

## AUTHOR INFORMATION

### Author Contributions

Dilara Diken has conducted the analysis and she has written the article. Dr. Mohit Mazumder and Elia Brodsky provided guidance throughout the project and directed the path taken for analysis. Dr. Harpreet Kaur has assisted with technical and analytical problems with the pipelines.

## ACKNOWLEDGMENTS

I thankfully acknowledge the team at Pine Biotech for their efforts on aiding this project. The groundwork of this project was conducted during the OmicsLogic Fellowship Program. The processing and visualization pipelines were prepared with the T-BioInfo server.

## ABBREVIATIONS

LET: Linear Energy Transfer
OTU: Operational Taxonomic Unit
NMDS: Non-metric Multidimensional Scaling PCoA: Principal Coordinate Analysis
PCA: Principal Component Analysis GABA: Gamma Aminobutyric Acid
GAD: Glutamic Acid Decarboxylase
GLS: Glutaminase
GS: Glutamine synthase
CNS: Central Nervous System RBE: Relative Biological Effects
ENS: Effective Number of Species

## Footnotes

* Template adapted from the American Chemical Society.

